# Female Anterior Cruciate Ligaments Exhibit a Muted Mechanobiological Response to Mechanical Loading

**DOI:** 10.1101/2025.05.16.654473

**Authors:** Lauren Paschall, Maxwell Konnaris, Erdem Tabdanov, Aman Dhawan, Spencer Szczesny

**Affiliations:** The Pennsylvania State University, Department of Biomedical Engineering, State College, PA; The National Institutes of Health, National Cancer Institute, Center for Cancer Research, Pediatric Oncology Branch, Bethesda, MD; The Pennsylvania State University, Department of Statistics, State College, PA; The Pennsylvania State University, Department of Orthopaedics and Rehabilitation, Hershey, PA; The Pennsylvania State University, Department of Pharmacology, Hershey, PA

**Keywords:** Mechanobiology, estrogen, anterior cruciate ligament, mechanical loading, tissue remodeling

## Abstract

Female athletes are significantly more likely to tear their anterior cruciate ligament (ACL) compared to their male counterparts. While there are several potential reasons for this, previous data from our lab demonstrated that female ACL explants have an impaired remodeling response to loading, which may prevent the repair of fatigue damage and lead to increased ACL rupture. The objective of this study was to identify the mechanisms driving the impaired remodeling of female ACLs to cyclic loading, including the role of estrogen. ACLs were harvested from male and female New Zealand white rabbits and cyclically loaded in a tensile bioreactor followed by bulk RNA-sequencing. Additional ACL explants treated with or without estradiol were analyzed using RT-qPCR to determine the regulatory effect of estrogen on markers for tissue remodeling and inflammatory cytokines with cyclic loading. We found that female ACLs exhibited significantly fewer differentially expressed genes (DEGs) in response to loading compared to male ACLs. Additionally, multiple mechanotransduction pathways were enriched with loading only in the male ACLs. While a few estrogen-related pathways were enriched in both male and female ACLs with loading, the expression of tissue remodeling markers was not different between estrogen treatment and vehicle control. Together, our findings highlight specific mechanotransduction pathways that may be responsible for the muted biological response of female ACLs to load, which provides a potential explanation for the increased rate of ACL tears in women.

## Introduction

While anterior cruciate ligament (ACL) tears are common in general during athletic activity^1^, female athletes are 2–8 times more likely to tear their ACLs compared to males^2–4^. Potential reasons for this include differences in anatomy, sex hormones, body size, and neuromuscular control^5–7^. However, women may also be more prone to ACL injury because of sex-specific ACL remodeling in response to fatigue loading. Recent studies suggest that ACL injuries occur from fatigue failure and accumulation of tissue damage^8^. Previous data from our lab demonstrated that female rabbit ACL explants have an impaired remodeling response to load. Specifically, we found that female ACLs fail to increase anabolic gene expression with loading and have increased catabolic gene expression compared to male ACL explants^8^, which may prevent the repair of fatigue damage explaining the increased rate of ACL tears in women. However, the cause of this sex difference in ACL mechanobiology is unknown.

Differences in sex hormones (estrogen) is one obvious potential reason for the sex-dependent ACL response to loading. Estrogen receptors (ERs) are present in both male and female ACL fibroblasts^9^. Additionally, estrogen influences multiple genes responsible for soft tissue remodeling^10–17^; however, the magnitude and directionality of this effect on the ACL are inconsistent^10,12,13,18–25^. Despite these discrepancies, clinical data demonstrate that estradiol concentrations correlate with increased knee laxity^26^, which alone is sufficient to explain an 8-fold increase in risk of ACL injury^27^. This effect can be attenuated through oral contraceptives that suppress spikes in estrogen levels^28,29^. Together, these data suggest that estrogen has a direct effect on ACL tissue mechanics and may contribute to ACL rupture.

Previous data also suggest that estrogen regulates the response of cells to mechanical loading. A prior study on ACL fibroblasts found that mechanical loading completely reverses the gene expression effect of estrogen^12^. However, the mechanisms underlying this interplay between estrogen and mechanical loading are unknown. In other tissues, ER signaling has been shown to interact with multiple mechanotransduction pathways (i.e., ERK, YAP/TAZ)^30–38^. However, the directionality of this effect is inconsistent. Specifically, studies on cancer cells show that ER signaling both increases^36–38^ and decreases^33–35^ YAP activation. While previous studies demonstrate that ER signaling regulates cellular mechanotransduction, it’s not possible to predict the effect of estrogen on ACL fibroblast mechanotransduction from prior data alone.

Therefore, the objective of this study was to identify potential mechanisms driving the impaired remodeling response of female ACLs. We conducted bulk RNA-sequencing on cyclically loaded male and female rabbit ACL explants to determine the innate sex differences in ACL mechanobiology in the native tissue environment. Additionally, we investigated the effect of estrogen treatment on the expression of remodeling genes in male and female loaded ACL explants. Our overall hypothesis was that female ACLs would exhibit an impaired mechanobiological response to load (increased catabolism, decreased anabolism) and this response will be influenced by estrogen. Specifically, we hypothesize that female ACLs will have altered mechanotransduction (suppression of ERK and YAP) as well as increased ER activity in response to load compared to males. We also hypothesized that estrogen treatment will increase catabolic and inflammatory gene expression in both male and female ACL explants in response to load.

## Materials and Methods

### ACL Harvest

A total of 33 male and female white New Zealand rabbits (2.8 – 3.2 kg, 14 – 16 weeks old) were euthanized (**Supplementary Table 1**) and ACLs were isolated under sterile conditions from a study approved by the Institutional Animal Care and Use Committee at Penn State University. ACLs were harvested following a previously published protocol^8^. Briefly, all surrounding tissues were removed, and the femur and tibia were cut into bone blocks to grip during mechanical loading. Cross-sectional area was determined by assuming an elliptical cross-section and major and minor diameters were determined using calipers.

### Mechanical Loading

ACLs were placed in a custom tensile bioreactor^39^ with culture media (low-glucose DMEM, 2% penicillin-streptomycin, 5% FBS, 25 mm HEPES, 4 mm GlutaMAX, 1 mm 2-Phospho-L-ascorbic acid trisodium) and kept at 37°C and 5% CO_2._ A subset of the ACLs was untreated to determine the innate mechanobiology of male and female ACL explants. To investigate the effect of estrogen, ACLs were treated with 10 nM 17β-estradiol (E2) (Sigma) or ethanol (0.1% v/v) as a vehicle control. The E2 concentration was based on a previous study investigating the role of estrogen and cyclic loading on isolated ACL fibroblasts^12^. Since underloading is detrimental to tendon/ligament homeostasis^40,41^ a 0.1 MPa static load was applied for 18 and 20 hours to acclimate the untreated and treated (vehicle/estrogen) ACLs to culture conditions respectively. The bioreactor then cyclically loaded the samples to 4 MPa at 0.5 hz for 8 h. This stress level was chosen since it represents the stress that exhibited the largest gene expression differences between male and female ACLs^8^. Control samples were maintained under a 0.1 MPa static load for the same duration.

### RNA Extraction

Immediately following loading, samples were removed and RNA was extracted following a previously published protocol^8^. Briefly, ACLs were excised from the bone blocks and were rinsed with ice cold RNase-free water and flash frozen in liquid nitrogen. The tissue was pulverized (6775 Freezer/Mill SPEX) and total RNA was extracted with RNeasy minicolumns (RNeasy Fibrous Tissue Kit, Qiagen). RNA concentration was quantified using a Qubit 4 Fluorometer (Thermo Fisher), and RNA integrity numbers (RINs) were determined using a TapeStation (Agilent).

### RNA-Sequencing

Samples that passed quality control (RIN > 6, total RNA > 250 ng) were sent to GENEWIZ (South Plainfield, NJ) for bulk RNA-Sequencing (**Supplementary Table 1**). cDNA libraries were prepared using the NEBNext Ultra II RNA Library Prep for Illumina using manufacturer’s instructions (NEB, Ipswich). The sequencing libraries were validated on the Agilent TapeStation (Agilent Technologies) and quantified using a Qubit 3.0 Fluorometer (Invitrogen) as well as qPCR (KAPA biosystems). The sequencing libraries were clustered on a lane of a NovaSeq 6000 S4 flow cell. The samples were sequenced with an average of 32 million reads per sample using a 2X150 bp paired end configuration. Image analysis and base calling were conducted by the Control software. Raw sequence data (.bcl files) generated by the sequencer were converted into fastq files and de-multiplexed using Illumina’s bcl2fastq 2.17 software. One mismatch was allowed for index sequence identification. Sequenced reads were trimmed to remove possible adapter sequences and nucleotides with poor quality using Trimmomatic v.0.36. The trimmed reads were mapped to the OryCun2.0 reference genome available on ENSEMBL using the STAR aligner v.2.5.2b. Unique gene hit counts were calculated by using FeatureCount from the Subread package v.1.5.2.

### Differential Gene Expression Analysis

Analysis and visualization of the bulk RNA-sequencing was performed utilizing the NIH Integrated Analysis Platform (NIDAP) bulk RNA-sequencing pipeline that utilizes R programs developed by a team of NCI bioinformaticians (Plantair Technologies, Denver, Colorado). Raw hit counts were uploaded and quality control measures were performed using principal component analysis and supervised hierarchical clustering to determine intergroup relationships and identify outliers. A female loaded sample (FL3) was identified as an outlier and removed (**Supplementary Fig. 1**). After outlier removal, normalization and DEG analysis was conducted utilizing the Limma Voom R package. Five comparisons were made to compare the response of each sex to load (female load, male load) and the relationships between sex in the fresh (sex effect fresh), static (sex effect static), and loaded (sex effect load) samples. The static control and male samples were used as the baseline for the comparisons. A moderated t-statistic was used to determine p-values, and a Benjamini-Hochberg false discovery rate was used to obtain the adjusted p-values.

Genes with an adjusted p-value < 0.05 and |log2 fold change| > 1 were identified as differentially expressed genes (DEGs). A list of DEGs were extracted from the model for each of the five comparisons.

### Pathway Analysis and Upstream Regulator Analysis

After differential gene expression analysis, all DEGs for each comparison were uploaded into Ingenuity Pathway Analysis (IPA, Qiagen, Version Q4 2024). Significantly enriched pathways were determined with a -log(p-value) > 1.3. Additionally, upstream regulator analysis was performed to determine potential regulators that could explain the observed changes in gene expression. To identify significantly activated or inhibited pathways or upstream regulators, a z-score was calculated in IPA for each pathway or upstream regulator. A positive z-score suggests inhibition while a positive z-score suggests activation. The magnitude of the z-score reflects the confidence of the predicted activation or inhibition state with |z-scores| > 2 considered strong prediction and |z-scores| > 1 considered moderate prediction.

### Reverse Transcription quantitative PCR (RT-qPCR)

The RNA extracted from each sample was diluted to 1 ng/ul, and cDNA was synthesized (High-Capacity cDNA Reverse Transcription Kit with RNase Inhibitor, Thermo Fisher). Quantitative polymerase chain reaction (qPCR) was performed using TaqMan probes (**Supplementary Table 2**) and a StepOne Plus Real-Time PCR system to measure the expression of anabolic (C*OL1A1*, *COL1A2*, *LOX, COL3A1, TGF*β*1, ACTA, TIMP1, TIMP3)*, catabolic (*MMP1, MMP2, MMP10, MMP13*), and inflammatory (*IL-1*β*, PTGS2)* genes. Additionally, ER target genes (*ESR1, GPER1, PGR*) were measured along with *GAPDH* as the reference gene. PCR efficiency and cycle number quantification (Cq) were obtained for each individual reaction using PCR-Miner (version 4.0)^42^. Samples that exhibited undetectable fluorescence for a given were excluded from the analysis for that gene. An outlier test (ROUT) was conducted to determine if any of the reaction efficiencies for each gene were an outlier. After removal of outliers, a single efficiency value was calculated for each probe by averaging all the individual sample efficiencies. Gene expression was quantified using the delta-delta Cq method correcting for primer efficiencies^43^.

The fold change for each sample was calculated as:

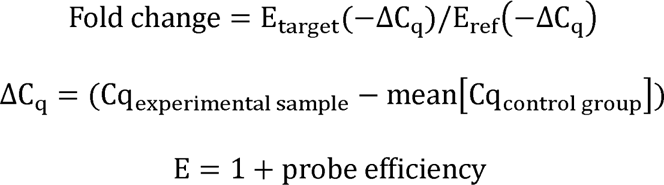

where the probe efficiencies are listed in **Supplementary Table 2**.

### Statistical Analysis of RT-qPCR Data

The fold change for each gene was log-normalized, and Mann-Whitney tests were conducted on the normalized data to compare the gene expression between treatments and sex. To determine the effect of mechanical loading on treatment, the estrogen treated and ethanol vehicle control samples were compared to their treatment-matched statically loaded control samples (for both males and females). To determine baseline treatment-specific gene expression differences, the estrogen treated static control samples were compared to the vehicle control static control samples (for both males and females). To determine baseline sex-specific gene expression differences, the female vehicle control samples were compared to the male vehicle control samples. Additionally, Mann-Whitney tests were used on the transformed data to determine the differential expression of each gene compared to the respective control condition. Given the numerous statistical comparisons, p-values were corrected using a false discovery rate analysis within each gene category (anabolic, catabolic, inflammatory, and ER target genes) with a Q-value of 5%. Statistical significance after corrections was set as *p* < 0.05 with statistical trends set at *p* < 0.10. All statistical analysis was performed using GraphPad Prism (version 10.3.1).

## Results

### Differential Gene Expression Analysis

In response to cyclic loading, male ACLs had 259 DEGs (122 upregulated, 137 downregulated) while female ACLs only had 10 DEGs (4 upregulated, 6 downregulated). When comparing the response to load between female and male ACLs (sex effect load), there are 116 DEGs (25 upregulated, 91 downregulated in female ACLs compared to males) (**Fig. 1A, Supplementary Tables 3 – 5)**. When looking at the top 20 DEGs in the male samples response to load, 75% of the DEGs were downregulated. Similarly, for the sex effect load comparison, 70% of the DEGs were further downregulated in female ACLs in response to load compared to males (**Table 1**). These differences are despite the fact that there are minimal differences between female and male ACLs prior to loading. Specifically, there were only 40 DEGs (19 upregulated, 21 downregulated) in female static ACLs compared to male samples, and only 2 DEGs (1 upregulated, 1 downregulated) in female freshly harvested ACLs compared to males (**Fig. 1B, Supplementary Table 6 - 7**).

**Figure 1:**
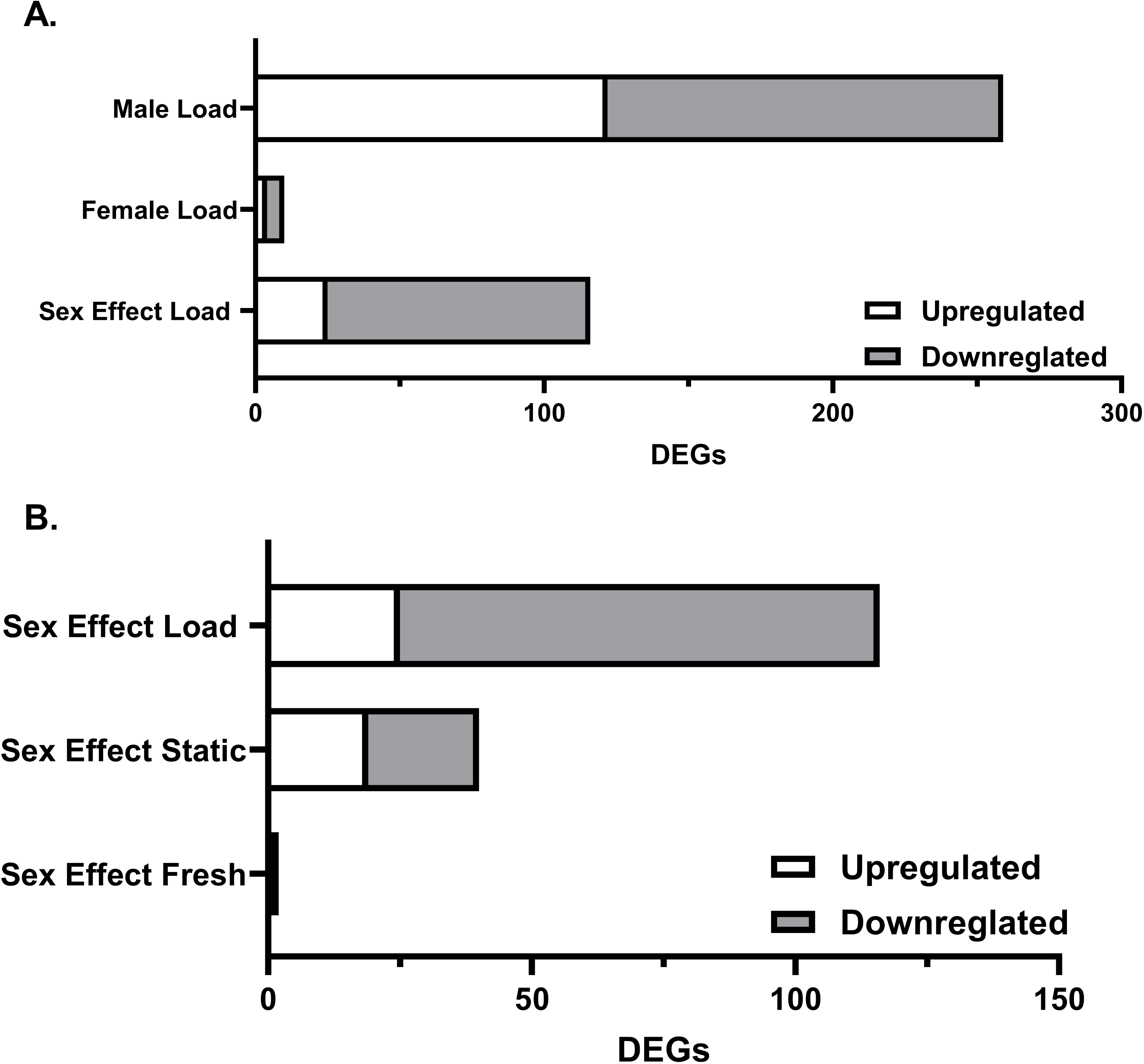
Differentially expressed genes. (A) Response of male and female ACLs in response to cyclic loading and comparing the response between sexes (sex effect load) (n = 3 males, n = 2 females). Male and female load are relative to their sex-matched static load. Sex effect load is comparing the female ACL response to load to the male ACL response to load. (B) Comparison of baseline gene expression differences of female ACLs relative to male ACLs (n = 3 static, n = 2 fresh). All DEGs had an absolute log2FC >1 and an adjusted p-value < 0.05.

**Table 1:**
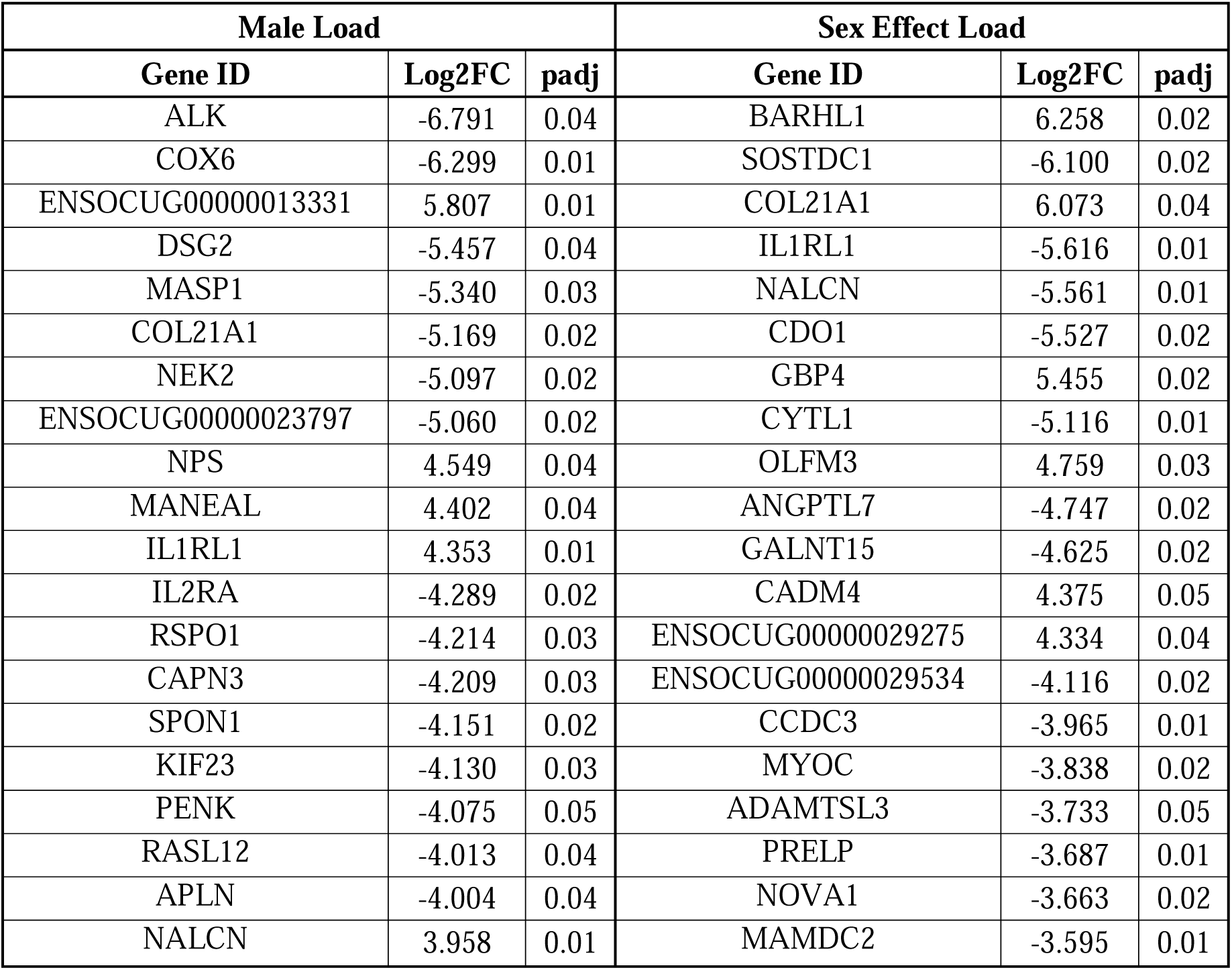
List of top 20 DEGs in response to load in male ACLs (and the sex effect of load (female ACLs response to load relative to male ACLs response to load).

Genes indicative of tissue remodeling (anabolic, catabolic, inflammatory markers)^8^ were searched in the list of DEGs for all five comparisons. In response to load, male ACLs downregulated catabolic gene *MMP2* and inflammatory marker *IL1*β while female ACLs exhibited no differentially expressed remodeling genes (**Table 2**). When comparing across sex, there are no differentially expressed remodeling genes for any of the comparisons. (**Table 2**).

**Table 2:** Differential expression of ECM remodeling genes in all comparisons. Bold text indicates statistical significance (p < 0.05). NAs indicate that the gene was not present in RNA-sequencing data.

| Gene Name | Male Load |  | Female Load |  | Sex Effect Load |  | Sex Effect Static |  | Sex Effect Fresh |  |
| --- | --- | --- | --- | --- | --- | --- | --- | --- | --- | --- |
|  | Log2FC | Padj | Log2FC | Padj | Log2FC | Padj | Log2FC | Padj | Log2FC | Padj |
| COL1A1 | 0.13 | 0.96 | -0.86 | 0.50 | -0.99 | 0.58 | 0.98 | 0.51 | 0.56 | 0.89 |
| COL1A2 | 0.26 | 0.85 | -0.91 | 0.36 | -1.17 | 0.40 | 0.95 | 0.40 | 1.22 | 0.60 |
| LOX | NA | NA | NA | NA | NA | NA | NA | NA | NA | NA |
| COL3A1 | -0.04 | 0.98 | -1.62 | 0.17 | -1.58 | 0.28 | 0.65 | 0.62 | 1.19 | 0.60 |
| TGFB1 | 0.06 | 0.92 | -0.06 | 0.91 | -0.12 | 0.85 | 0.10 | 0.86 | -0.08 | 0.95 |
| ACTA2 | -2.40 | 0.11 | -0.22 | 0.88 | 2.18 | 0.24 | -0.72 | 0.59 | -1.22 | 0.61 |
| TIMP1 | 0.55 | 0.47 | -0.54 | 0.40 | -1.09 | 0.25 | 0.12 | 0.92 | -0.62 | 0.59 |
| TIMP3 | NA | NA | NA | NA | NA | NA | NA | NA | NA | NA |
| MMP1 | -2.27 | 0.11 | 0.96 | 0.49 | 3.23 | 0.12 | -1.12 | 0.49 | 0.38 | 0.90 |
| MMP2 | -1.62 | **0.03** | -0.27 | 0.66 | 1.36 | 0.14 | -0.82 | 0.23 | 0.29 | 0.85 |
| MMP10 | NA | NA | NA | NA | NA | NA | NA | NA | NA | NA |
| MMP13 | NA | NA | NA | NA | NA | NA | NA | NA | NA | NA |
| IL1B | -2.13 | **0.06** | -0.53 | 0.56 | 1.60 | 0.25 | -1.22 | 0.23 | -0.65 | 0.94 |
| PTGS2 | NA | NA | NA | NA | NA | NA | NA | NA | NA | NA |

### Differentially Expressed ER Target Genes

Previously reported ER target genes^44,45^ were searched in the list of DEGs for all five comparisons. In response to load, male ACLs upregulated *ESR1* and multiple genes trended towards downregulation (*IGFBP4, TPD52L1, SERPINE1*) while female ACLs exhibited no differentially expressed ER target genes (**Table 3**). When comparing the DEGs across sex, there were surprisingly no differentially expressed ER target genes for the freshly harvested or static samples. In response to load, *ESR1* trended towards downregulation in female loaded ACLs compared to male loaded ACLs (**Table 3**).

**Table 3:** Differential expression of estrogen receptor target genes in all comparisons. Bold text indicates statistical significance (p < 0.05) and italicized text indicates trends (p < 0.10).

|  | Male Load |  | Female Load |  | Sex Effect Load |  | Sex Effect Static |  | Sex Effect Fresh |  |
| --- | --- | --- | --- | --- | --- | --- | --- | --- | --- | --- |
| Gene Name | Log2FC | Padj | Log2FC | Padj | Log2FC | Padj | Log2FC | Padj | Log2FC | Padj |
| ESR1 | 3.542 | **0.02** | -0.662 | 0.61 | -4.204 | **0.05** | 2.386 | 0.16 | 0.339 | 0.81 |
| IGFBP4 | -1.989 | *0.10* | -0.750 | 0.49 | 1.238 | 0.42 | -0.583 | 0.62 | -1.360 | 0.52 |
| TPD52L1 | 2.396 | *0.10* | -1.047 | 0.61 | -3.444 | 0.17 | 0.158 | 0.96 | 1.560 | 0.49 |
| GPER1 | -2.192 | 0.12 | 0.360 | 0.75 | 2.552 | 0.16 | -0.620 | 0.59 | -0.213 | 0.96 |
| STC2 | -1.870 | 0.35 | 1.350 | 0.37 | 3.219 | 0.20 | -0.188 | 0.95 | 0.348 | 0.95 |
| RBBP8 | -0.212 | 0.70 | 0.677 | 0.16 | 0.889 | 0.17 | -0.031 | 0.97 | -0.097 | 0.97 |
| SIAH2 | 0.944 | 0.18 | -0.025 | 0.98 | -0.968 | 0.29 | 0.587 | 0.41 | 1.010 | 0.40 |
| ADCY9 | 0.249 | 0.70 | -0.201 | 0.76 | -0.450 | 0.56 | -0.020 | 0.98 | -0.661 | 0.45 |
| SERPINE1 | -1.830 | *0.08* | 0.259 | 0.82 | 2.089 | 0.15 | -0.706 | 0.55 | 0.810 | 0.61 |
| EPHA4 | -1.150 | 0.29 | -0.348 | 0.73 | 0.802 | 0.55 | -0.059 | 0.97 | -0.906 | 0.60 |

### Pathway Analysis of ACLs Response to Cyclic Load

In response to cyclic loading, IPA found that male ACLs had 139 signaling pathways enriched, while female ACLs only had 31 enriched pathways (**Supplementary Table 8 – 9**). When directly comparing across sex (sex effect load), there were 82 pathways enriched (**Supplementary Table 10**).

We then investigated which estrogen-related and mechanotransduction pathways were enriched with loading. For estrogen, extra-nuclear estrogen signaling was enriched in male ACLs in response to load and was trending towards inhibition (z-score: -1), while ESR-mediated signaling was enriched in female ACLs in response to load (**Supplementary Table 8 – 9**). For mechanotransduction, male ACLs had 15 pathways enriched in response to load (**Table 4**) with 8 pathways predicted to be downregulated and zero pathways predicted to be upregulated (**Table 4**). The female ACLs exhibited no enriched pathways. Additionally, there were 9 pathways enriched when directly comparing across sex (sex effect load). Prominent pathways of interest included integrin signaling, FAK signaling, MAPK signaling, PI3K/AKT signaling, Rho GTPase cycle, and G-protein coupled receptor signaling. When comparing enriched pathways between male and female ACLs in response to load, 5 pathways were enriched in both male load and sex effect of load, which were generally predicted to be inhibited (**Table 5**). Interestingly, FAK signaling was predicted to be inhibited in the sex effect load comparison, indicating an increased inhibition of FAK in female ACLs compared to male ACLs in response to load.

**Table 4:**
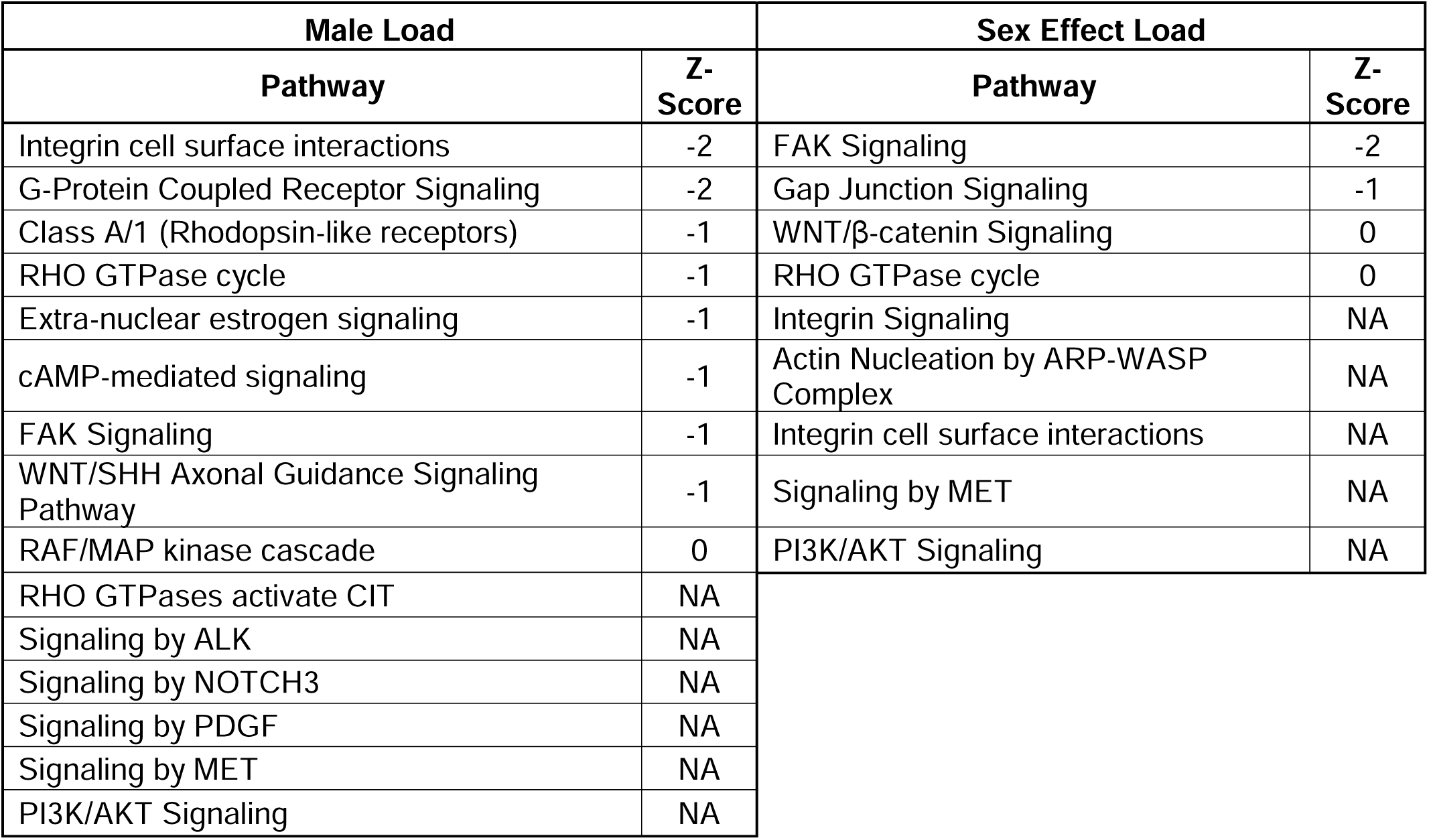
Mechanotransduction pathways enriched in response to load. Pathways with a z-score > 0 are predicted to be activated and z-score < 0 are predicted to be inhibited.

**Table 5:** Mechanotransduction pathways enriched in both male ACL response to load and sex effect load. Pathways with a z-score > 0 are predicted to be activated and z-score < 0 are predicted to be inhibited.

| Pathway | Male Load Z-Score | Sex Effect Load Z-Score |
| --- | --- | --- |
| FAK Signaling | -1 | -2 |
| Integrin cell surface interactions | -2 | NA |
| RHO GTPase cycle | -1 | 0 |
| Signaling by MET | NA | NA |
| PI3K/AKT Signaling | NA | NA |

Finally, we also investigated pathways associated with extracellular matrix (ECM) organization. Male ACLs had 8 ECM pathways enriched in response to load (**Table 6**), while female ACLs had no enriched pathways. Three pathways were enriched when directly comparing across sex (sex effect load). These included collagen synthesis and degradation, activation and inhibition of matrix metalloproteinases, and elastic fiber formation. Interestingly, most of these pathways were predicted to be inhibited in the male ACLs response to load. When looking at what pathways are different between male ACLs and female ACLs in response to load, 2 pathways related to collagen synthesis were enriched in both the male load and sex effect of load comparisons (**Table 7**).

**Table 6:** Extracellular matrix pathways enriched in response to load. Pathways with a z-score > 0 are predicted to be activated and z-score < 0 are predicted to be inhibited.

| Male Load |  | Sex Effect Load |  |
| --- | --- | --- | --- |
| Pathway | Z-Score | Pathway | Z-Score |
| Collagen degradation | -2 | Elastic fibre formation | NA |
| Inhibition of Matrix Metalloproteases | 2 | Collagen biosynthesis and modifying enzymes | NA |
| Degradation of the extracellular matrix | -2 | Collagen chain trimerization | NA |
| Extracellular matrix organization | -2 |  |  |
| Collagen biosynthesis and modifying enzymes | -2 |  |  |
| Collagen chain trimerization | -2 |  |  |
| Assembly of collagen fibrils and other multimeric structures | NA |  |  |
| Activation of Matrix Metalloproteinases | NA |  |  |

**Table 7:**
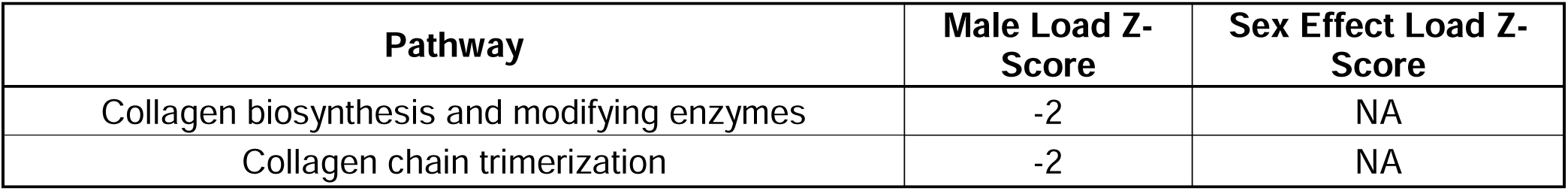
Extracellular matrix pathways enriched in both male ACLs response to load and sex effect load. Pathways with a z-score > 0 are predicted to be activated and z-score < 0 are predicted to be inhibited.

### Upstream Regulator Analysis of ACLs Response to Cyclic Load

IPA upstream regulator analysis identified several key regulators of male ACL response to load (**Table 8**), including multiple regulators related to sex hormones (beta-estradiol, *ERBB2*, dihydrotestosterone, *AR, ESR1*), growth factors (TGFβ1, VEGF, IGF1) and inflammatory cytokines (TNF, LPS, IL1β, prostaglandin E2), with the majority of these regulators predicted to be inhibited. There were no significant upstream regualtors in the female ACLs response to load (**Table 8**). In the sex effect load, *AR* and TGFβ1 were predicted to be inhibited while TNF was predicted to be activated.

**Table 8:**
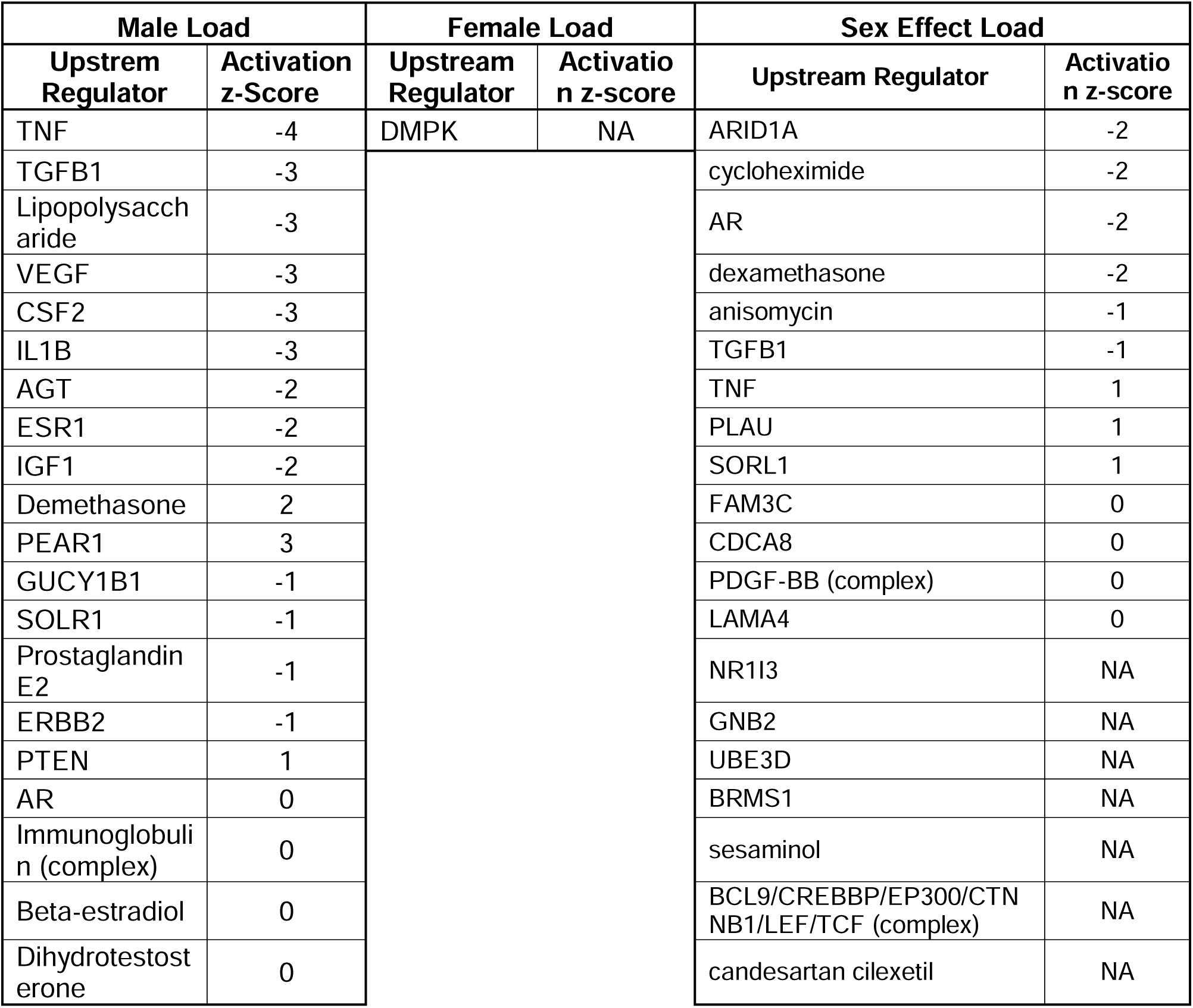
List of top 20 predicted upstream regulators of response to load in female and male ACLs (relative to sex-matched static control) and the sex effect of load (female ACL response to load relative to male ACL response to load). Regulators with a z-score > 0 are predicted to be activated and z-score < 0 are predicted to be inhibited.

### Effects of Estrogen Treatment on the Remodeling Response of ACLs to Cyclic Load

When comparing the expression of remodeling genes between male and female statically loaded ACLs cultured without estrogen (vehicle control), there were multiple anabolic (*COL1A2, LOX, TGF*β*1, TIMP3*), catabolic (*MMP1, MMP2, MMP13*), and inflammatory (*IL1*β*, PTGS2*) markers significantly upregulated in the female ACLs compared to males and *MMP1*0 was trending toward upregulation (**Supplementary Fig. 2A**). However, there was no differential expression of any of the anabolic, catabolic, or inflammatory markers with estrogen treatment in the statically loaded male or female ACLs (**Fig. 2 A-B**). This was also true when comparing the response of estrogen treatment between male and female statically loaded ACLs (**Fig. 2C**).

**Figure 2:**
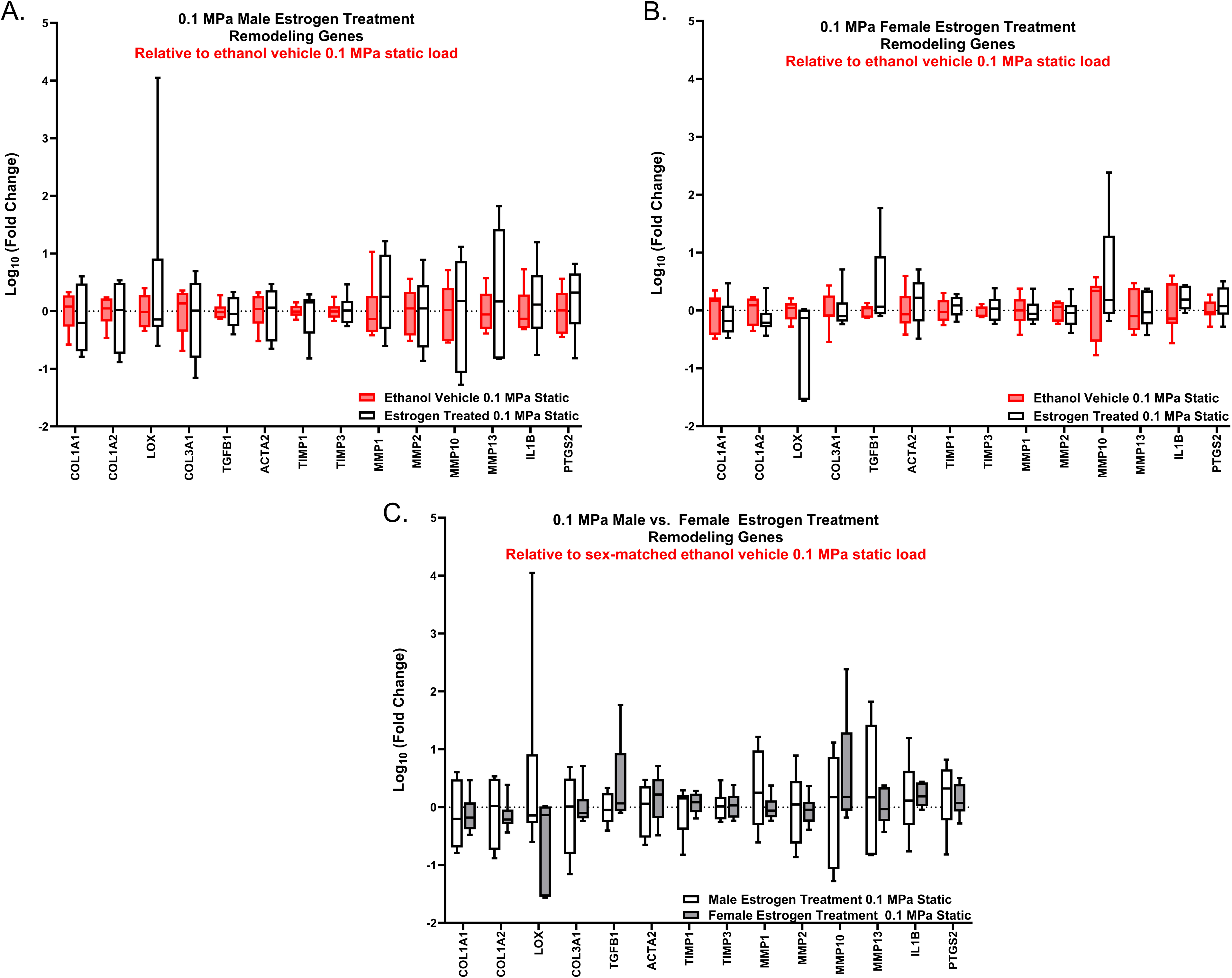
Basal remodeling gene expression response of estrogen treatment. RT-qPCR analysis of statically loaded estrogen treated ACLs relative to their sex-matched ethanol vehicle control (represented by the red box and whisker plots). (A) Statically loaded male ACL response. (B) Statically loaded female ACL response. (C) Comparison of statically loaded male and female ACL response to estrogen treatment (n = 6 for all male samples, n =7 for female vehicle, n = 5 – 6 for female estrogen treated). Data represented as box and whisker plot with the whiskers representing the min and max data.

No differential gene expression was observed with cyclical loading in the male ACLs compared to static controls with vehicle or estrogen treatment (**Fig. 3 A-B**). Direct comparisons between the vehicle control and estrogen-treated male samples found that there was no effect of estrogen on gene expression (**Supplementary Fig. 3A**). Consistently, the female cyclically loaded ACLs exhibited no differential gene expression compared to static controls with either vehicle or estrogen treatment (**Fig. 3 C-D**). Direct comparisons between vehicle and estrogen-treated cyclically loaded female samples found no effect of estrogen on gene expression (**Supplementary Fig. 3B**). Finally, direct comparison between the male and female cyclically loaded ACLs found that there were no sex differences in the vehicle or estrogen-treated groups (**Supplementary Fig. 3 C-D**).

**Figure 3:**
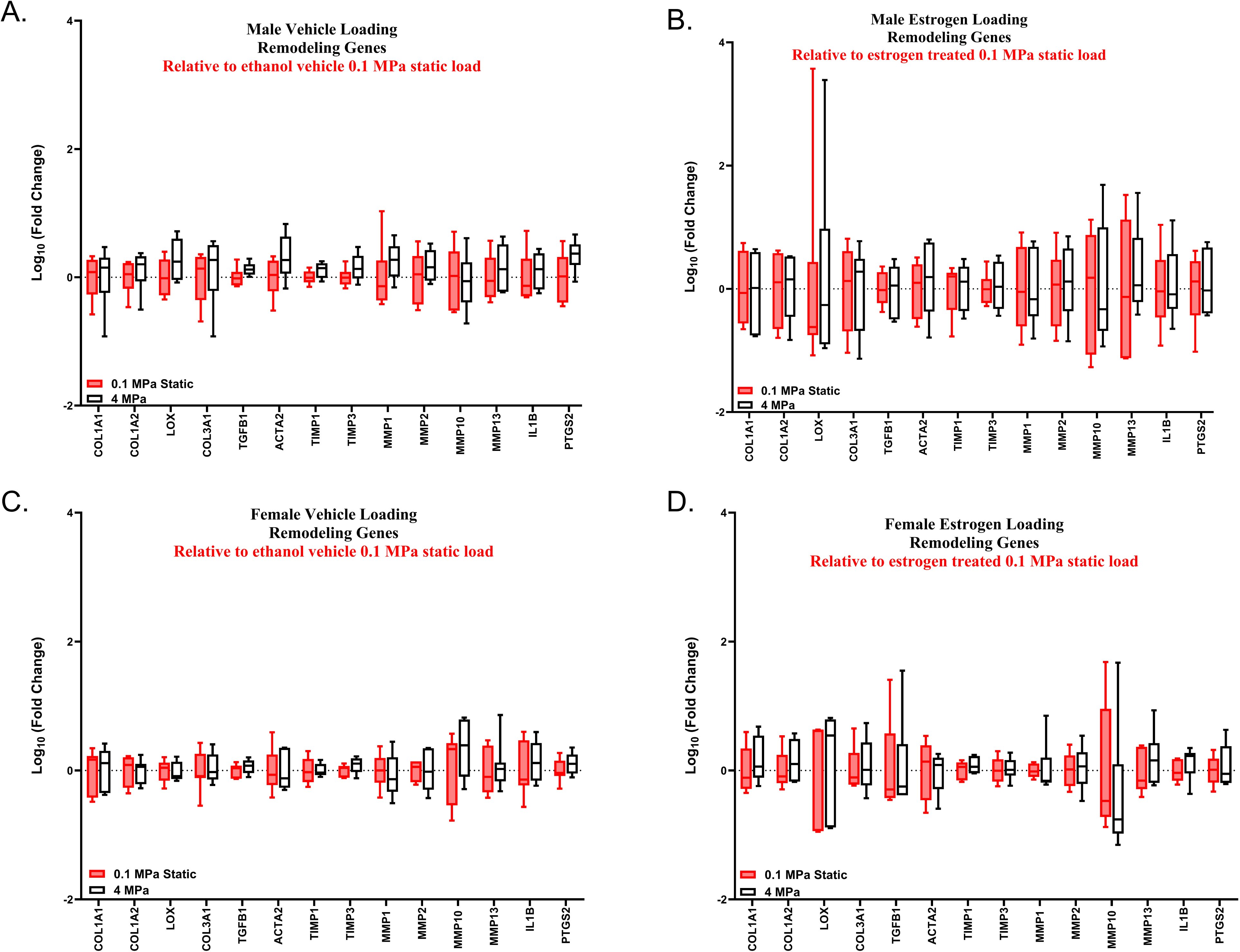
Effect of estrogen treatment on ACL remodeling in response to load. RT-qPCR analysis of cyclically loaded male and female ACLs relative to their sex and treatment-matched 0.1 MPa static load (represented by the red box and whisker plots). (A) Gene expression changes in vehicle control male ACLs. (B) Gene expression changes in estrogen treated male ACLs. (C) Gene expression changes in vehicle control female ACLs. (D) Gene expression changes in female estrogen treated ACLs (n = 6 for all male samples, n =7 for female vehicle, n = 6 for female estrogen treated). Data represented as box and whisker plot with the whiskers representing the min and max data.

### Effects of Estrogen Treatment on Estrogen Target Genes of ACLs to Cyclic Load

In contrast to our RNA-sequencing data (sex effect static), we found *GPER1* to be significantly upregulated in the statically loaded female ACLs compared to males without estrogen treatment (**Supplementary Fig. 2B**). However, there was no differential expression of any of the ER genes in the statically loaded male or female ACLs in response to estrogen treatment (**Fig. 4 A-B**). Additionally, there were no differences in gene expression when comparing the response of estrogen treatment between male and female statically loaded ACLs (**Fig. 4C**).

**Figure 4:**
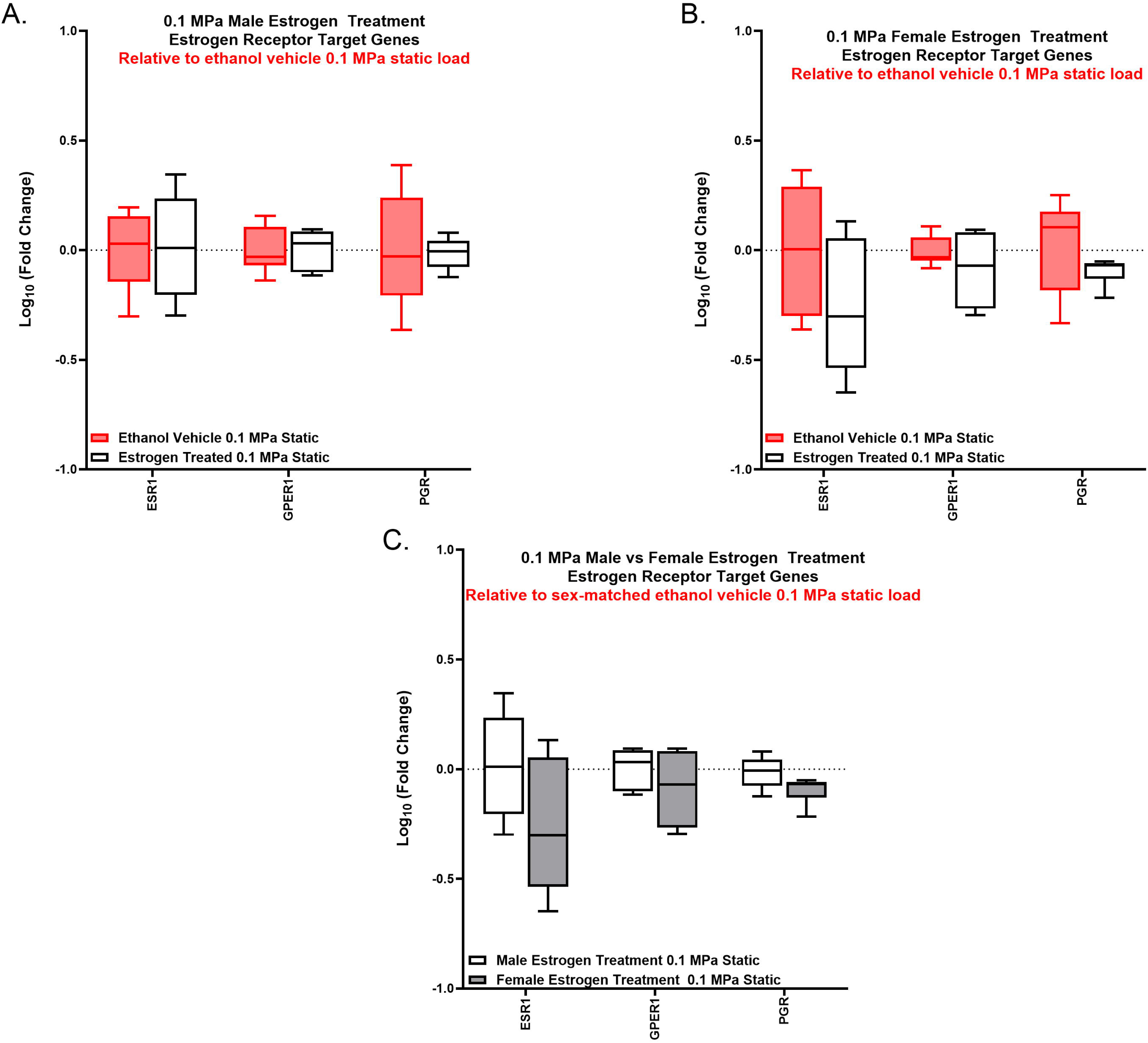
Basal estrogen receptor target genes response to estrogen treatment. RT-qPCR analysis of statically loaded estrogen treated ACLs relative to their sex-matched ethanol vehicle control (represented by the red box and whisker plots). (A) Statically loaded male ACL response. (B) Statically loaded female ACL response. (C) Comparison of statically loaded male and female ACLs response to estrogen treatment (n = 5 for male vehicle, n = 6 for male estrogen treated, n = 6 - 7 for female vehicle, n = 6 for female estrogen treated). Data represented as box and whisker plot with the whiskers representing the min and max data.

Consistent with our findings with the markers of tissue remodeling, there was no differential expression of the estrogen target genes with cyclic loading in the male (**Fig. 5 A-B**) or female samples (**Fig. 5 C-D**) with or without estrogen treatment. Similarly, there was no effect of estrogen (**Supplementary Fig. 4A-B**) or sex (**Supplementary Fig. 4C-D**) on the expression of estrogen target genes in response to loading.

**Figure 5:**
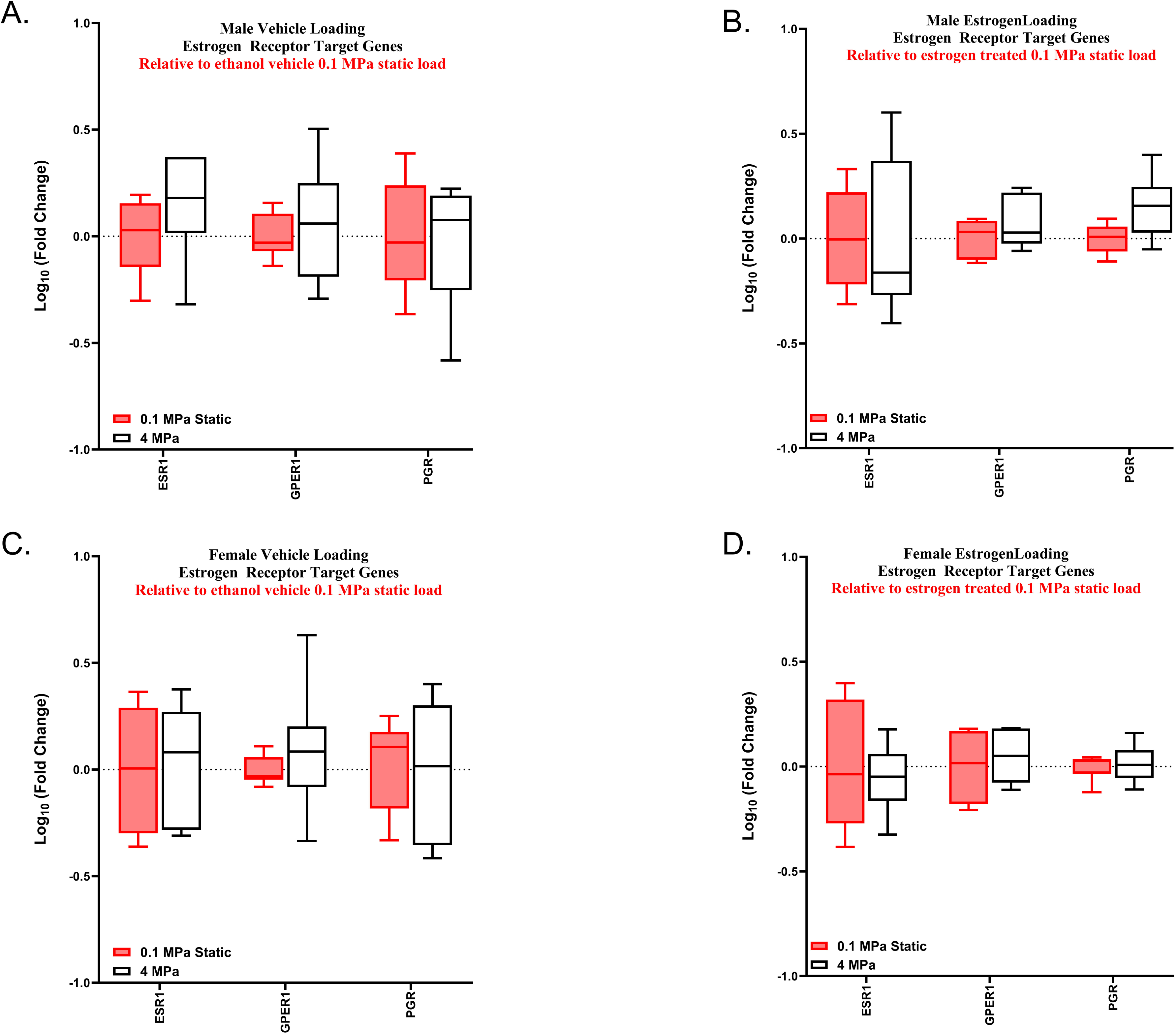
Effect of estrogen treatment on estrogen receptor target genes in response to load. RT-qPCR analysis of cyclically loaded male and female ACLs relative to their sex and treatment-matched 0.1 MPa static load. (A) Gene expression changes in vehicle control male ACLs. (B) Gene expression changes in estrogen treated male ACLs. (C) Gene expression changes in vehicle control female ACLs. (D) Gene expression changes in female estrogen treated ACLs (n = 6 for all male samples, n =6 - 7 for female vehicle, n = 6 for female estrogen treated). Data represented as box and whisker plot with the whiskers representing the min and max data.

## Discussion

The objective of this work was to investigate potential mechanisms driving the impaired remodeling response of female ACLs in response to load. Consistent with our hypothesis, our RNA-sequencing data demonstrated that female ACLs had a muted response to mechanical loading compared to male ACLs. Specifically, female ACLs had significantly fewer DEGs in response to load compared to male ACLs (**Fig. 1A**). This is despite the fact that there were minimal differences in gene expression between freshly harvested female and male ACLs (**Fig. 1B**). Suggesting, that the differences in the number of DEGs in response to load between male and female ACLs are due to differences in mechanobiology rather than basal biological differences. Furthermore, multiple mechanotransduction pathways were significantly enriched in male ACLs in response to load while none were enriched in female ACLs (**Table 4**). Surprisingly, half of these mechanotransduction pathways were predicted to be downregulated in the male samples. However, a number of prominent mechanotransduction pathways, including MAPK (i.e., ERK) and PI3K/AKT, were enriched without likely being downregulated, suggesting that these pathways may be important for the mechanobiological response of male ACLs. Additionally, when comparing the response to load between male and female ACLs, FAK (focal adhesion kinase) signaling was predicted to be further inhibited with load in female ACLs compared to male ACLs (**Table 4, 5**). Therefore, FAK signaling may also help explain sex differences in ACL mechanobiology given that FAK plays a prominent role in cell-matrix mechanotransduction and regulates the mechanoresponse of tendon cells^46^. Together, our RNA-sequencing data support the hypothesis that female ACLs have a muted mechanobiological response to load potentially inhibiting their ability to repair tissue damage that predisposes the ACL to rupture.

Our secondary hypothesis was that the impaired mechanobiological response in female ACLs is due to estrogen signaling. Surprisingly, there were no basal differences in ER target genes between male and female ACLs (**Table 3**). This could be due to the fact that we did not measure estrogen concentrations to confirm that estrogen levels were higher in female rabbits at the time of euthanasia. However, since rabbits are induced ovulators, female rabbits exhibit relatively stable serum estrogen concentrations over time^47^. Additionally, previous studies suggest that serum estrogen concentration is 10 times higher in female rabbits (20 – 50 pg/ml)^19,20^ compared to male rabbits (3 pg/ml)^48^. These data suggest that the estrogen concentration should be consistently higher in the female ACL samples, suggesting that the chosen estrogen targets are not representative of estrogen signaling in rabbits. Downstream targets of estrogen activation are tissue and species specific^49,50^, and there are no data available on ER target genes in the ACL or rabbit, providing difficulties in validating ER activation. Additionally, we did not observe differential expression of the chosen ER targets in the female ACLs in response to load. However, the ESR-mediated signaling pathway was enriched in the female ACL response to load (**Supplementary Table 9**). Additionally, the extra-nuclear estrogen signaling was enriched and trending toward inhibition in the male ACL response to load (**Supplementary Table 8**). Furthermore, upstream regulator analysis revealed inhibition of ESR1 in the male ACL response to load (**Table 8**). Therefore, while rabbit-specific markers of estrogen signaling may need to be identified, our RNA-sequencing data suggest that estrogen-related signaling may help explain sex differences in ACL mechanobiology.

To further investigate this, we cyclically loaded male and female ACLs in the presence of exogenous estradiol and measured gene expression of markers of ECM remodeling. Unfortunately, we weren’t to replicate our previous work that utilized the identical loading protocol and identified a sex dependent response in these genes with loading^8^. Given that there was no effect of cyclic loading on our chosen remodeling genes, we were unable to evaluate the effect of estrogen (**Fig. 3, Supplementary Fig. 3 A-B**). It’s possible that the lack of effect with cyclic loading is due to a poor choice of gene markers for tissue remodeling. Specifically, while our RNA-sequencing data found 259 DEGs with loading in male ACLs, only two of those genes were chosen as markers of tissue remodeling for our PCR experiments (*MMP2*, *IL1*β) (**Table 2**). However, when looking at the pathway analysis related to extracellular organization, there were only three pathways enriched when comparing the response to load between female and male ACLs (**Table 6**). Furthermore, only a few genes enriched in these pathways are relevant to remodeling of the bulk ECM (MMP2, MMP14, COL5A3), whereas the others are more relevant to the local pericellular matrix (ADAM10, ADAM12, COL4A1, COL4A2, LAMB1) (**Supplementary Table 11**). Together, this suggests that the mechanobiological difference between male and female ACLs may not involve bulk ECM remodeling. Still, the RNA-sequencing data suggest that ACLs do have a sex-dependent mechanobiological response, and future work is needed to identify the role of estrogen in driving this differential response.

It’s also possible that the absence of an effect of estrogen on ACL explant gene expression was due to insufficient estradiol treatment. However, we used the same concentration (10^-8^ M) that increased expression of *COL1* and *COL3* in a previous study of isolated ACL fibroblasts^12^. Additionally, our chosen concentration is three orders of magnitudes greater than serum estrogen concentrations found in rabbits (3.5 x 10^-11^ M)^19,20^. Another possibility is that there was sufficient estradiol in the culture media to saturate estrogen signaling. While we did not use charcoal-stripped FBS in our culture media, there is only approximately 2 pg/ml (< 10^-11^ M) of estradiol in 10% FBS, which was not sufficient to saturate the response of isolated ACL fibroblasts to estradiol^12^. A final possibility is that the effect of estrogen is no longer present at the end of our culture period (28 h), given that the effect of estrogen treatment on collagen gene expression decreased after 24 h^12^.

A final limitation to this study was that our RNA-sequencing sample size was small for the loaded female samples due to the removal of an outlier (n=2) (**Supplementary Fig. S1**). While it’s possible that having a smaller sample size could reduce statistical power and lead to fewer DEGs, this is likely not the reason for the low number of DEGs in the female ACLs response to load. Specifically, it’s clear from the PCA plot of the samples included in the sequencing analysis **(Supplementary Fig. S1B**) that the male loaded samples separated from the male static samples to a greater amount than the female samples, which is consistent with the fewer number of DEGs in the female ACLs in response to load.

In conclusion, our findings suggest that female ACLs have a muted mechanobiological response to load compared to male ACLs. This may be due to decreased activation of mechanotransduction pathways, including FAK, MAPK (i.e., ERK), and PI3K/AKT, as well as increased estrogen signaling in female ACLs in response to load. These data support the novel hypothesis that females exhibit an impaired mechanobiological response to load making female athletes more susceptible to ACL tears. Future work could compare the structural and mechanical changes that occur between male and female ACLs in response to load. This will enable a direct comparison of male and female ACLs ability to repair tissue damage. Future work is needed to better understand the role of estrogen signaling in ACL mechanobiology. Together, these data can provide insight into the ability of female ACLs to repair tissue damage and possible approaches for reducing the risk of ACL ruptures in females by modulating estrogen levels (e.g., contraceptive use).

## Acknowledgements

We would like to acknowledge the Genomics Core at Penn State for running the PCR plates. Funding for this project was provided by the Orthopaedic Research and Education Foundation (234995) and the Congressionally Directed Medical Research Program (W81XWH2110152).

## Figure Captions

**Supplementary Figure 1:**
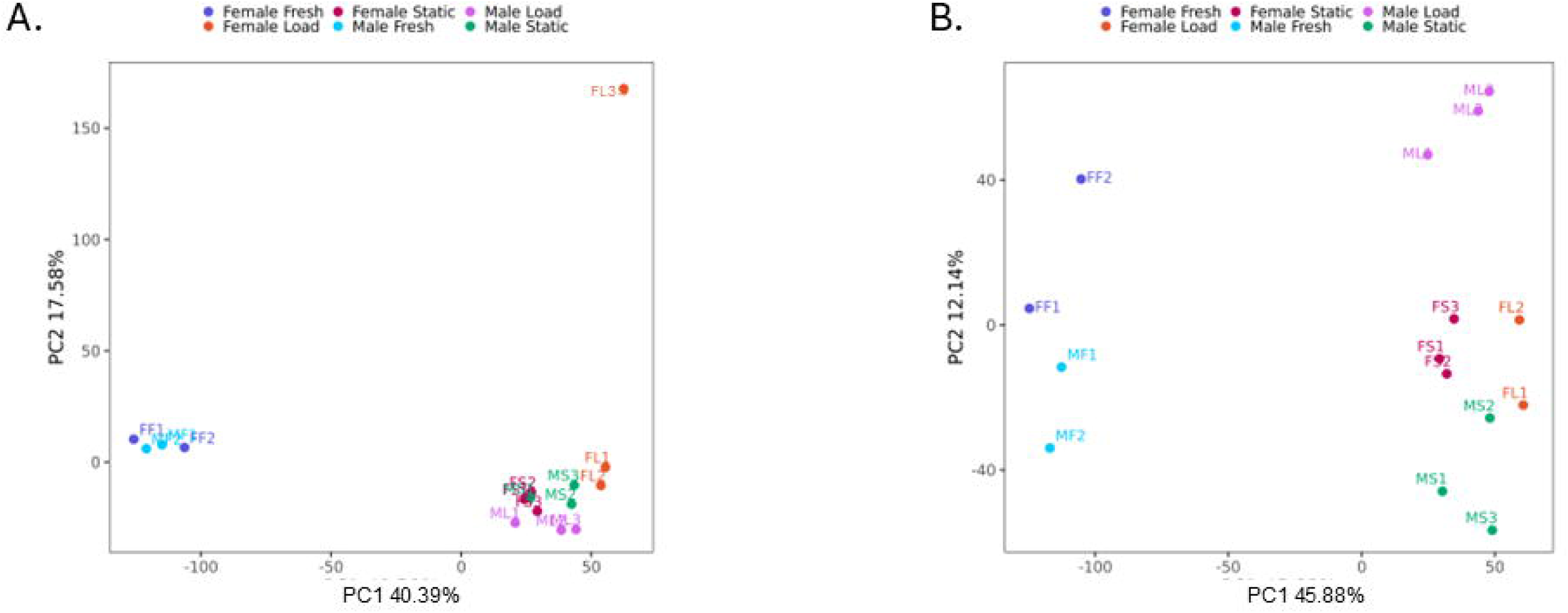
Principal component analysis. Visual distribution of samples based on two principal components. Each point represents an individual sample, labeled with the sample number. (A) Principal component analysis without the outlier removal (FL3). (B) Principal component analysis with the outlier removal.

**Supplementary Figure 2:**
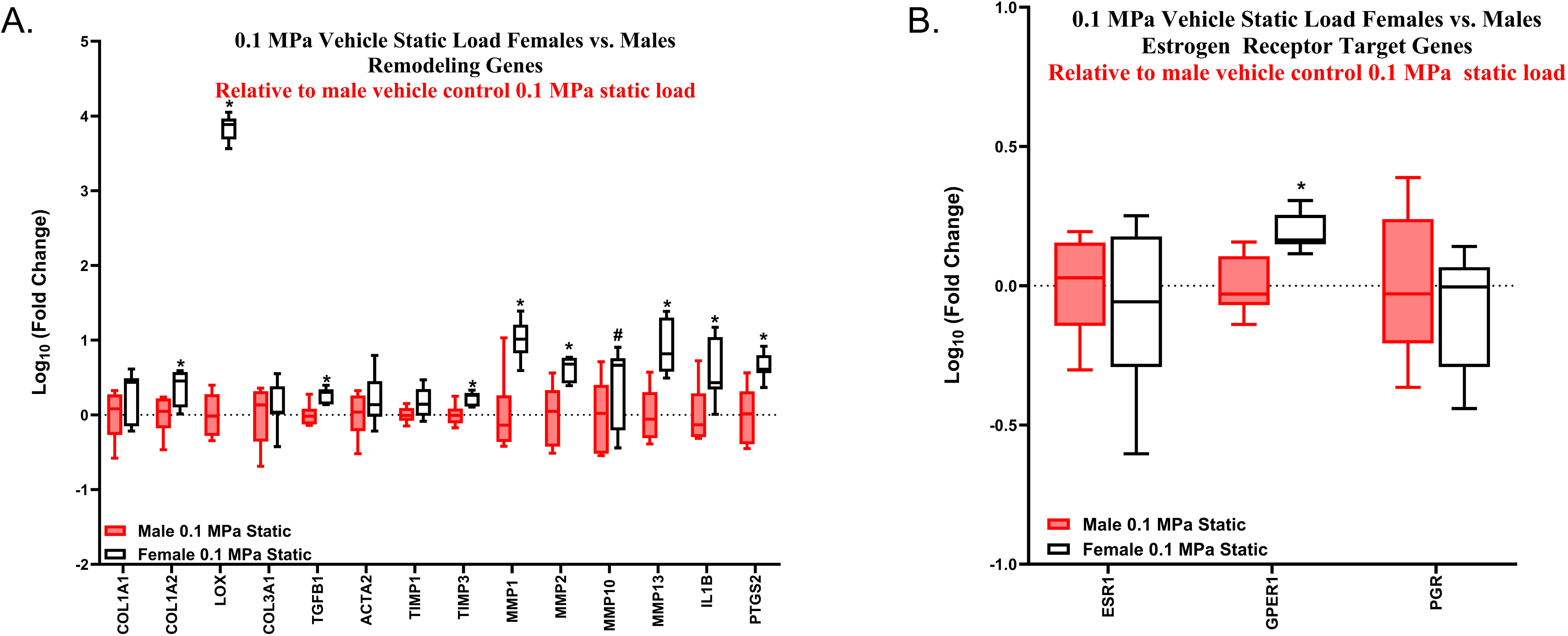
Comparison of vehicle control female and male ACL gene expression at baseline. RT-qPCR analysis of statically loaded female samples relative to male statically loaded samples. (A) Remodeling genes. (B) Estrogen receptor target genes (n = 6 for male samples and n = 7 for female samples for remodeling genes, n = 6 for male samples and n = 6-7 for female samples for estrogen receptor target genes) *p < 0.05, #p < 0.1. Data represented as box and whiskers plot with the whiskers representing the min and max data.

**Supplementary Figure 3:**
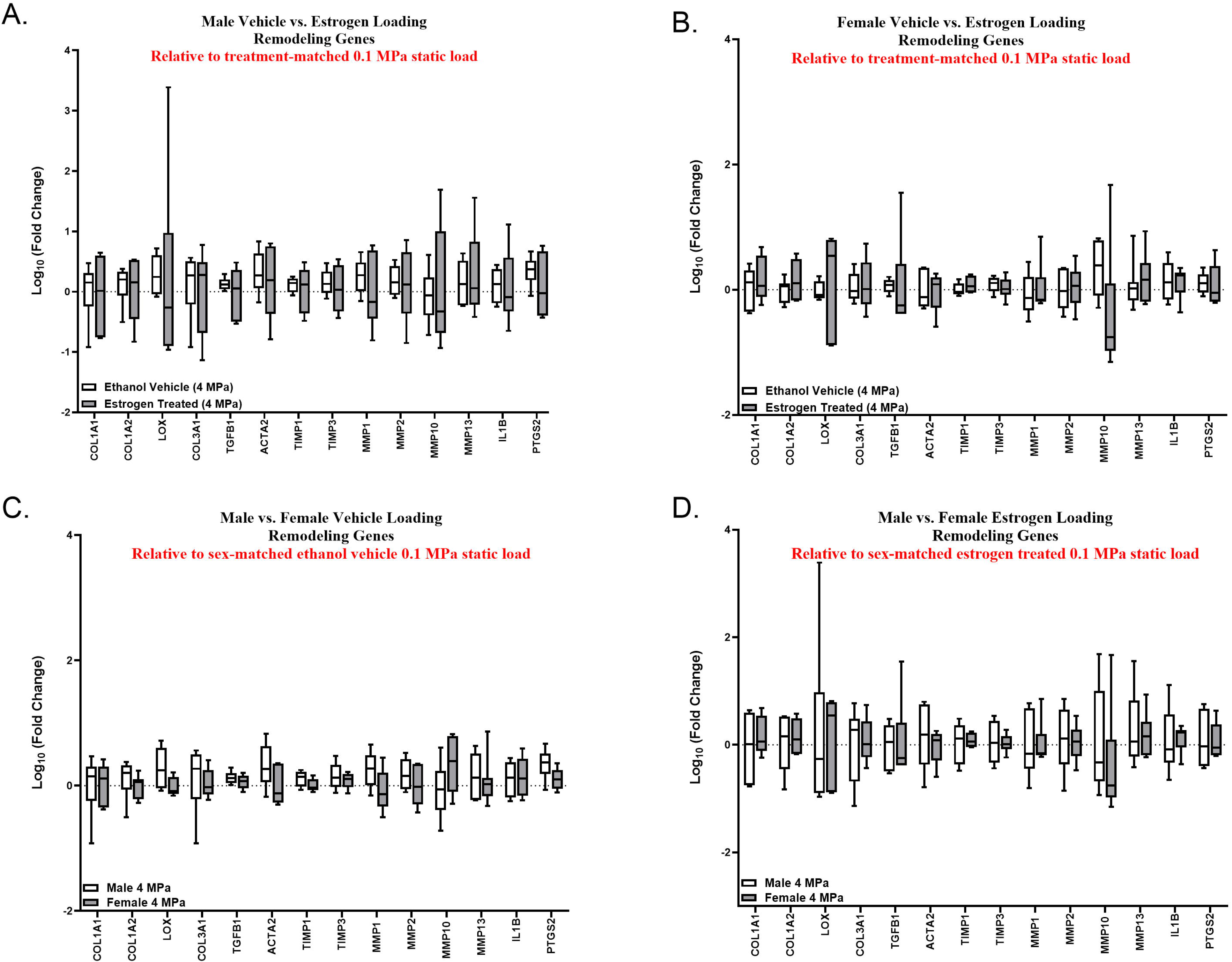
Comparisons of the male and female ACLs remodeling response to load. RT-qPCR analysis of cyclically loaded male and female ACLs relative to their sex and treatment-matched 0.1 MPa static load. (A) Comparison of vehicle control and estrogen treatment in male samples. (B) Comparison of vehicle control and estrogen treatment in female samples. (C) Comparison of male and female ACLs response to vehicle treatment. (D) Comparison of male and female ACLs response to estrogen treatment (n = 6 for all male samples, n =7 for female vehicle, n = 6 for female estrogen treated). Data represented as box and whisker plot with the whiskers representing the min and max data.

**Supplementary Figure 4:**
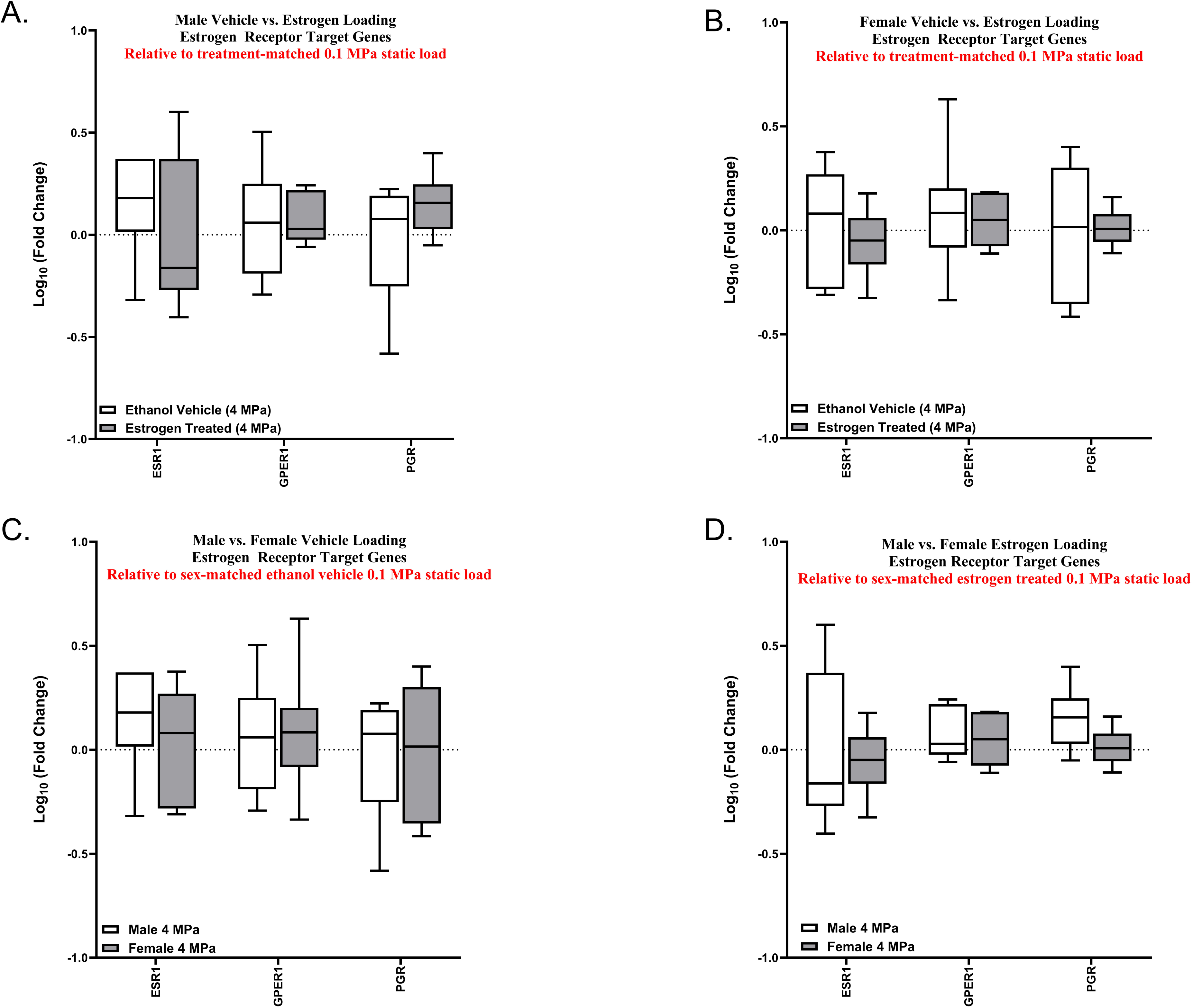
Comparisons of the male and female ACLs estrogen receptor target genes in response to load. RT-qPCR analysis of cyclically loaded male and female ACLs relative to their sex and treatment-matched 0.1 MPa static load. (A) Comparison of vehicle control and estrogen treatment in male samples. (B) Comparison of vehicle control and estrogen treatment in female samples. (C) Comparison of male and female ACL response to vehicle treatment. (D) Comparison of male and female ACLs response to estrogen treatment (n = 6 for all male samples, n =6 - 7 for female vehicle, n = 6 for female estrogen treated). Data represented as box and whisker plot with the whiskers representing the min and max data.

